# Sensory Entrained TMS (seTMS) enhances motor cortex excitability

**DOI:** 10.1101/2024.11.26.625537

**Authors:** Jessica M. Ross, Lily Forman, Juha Gogulski, Umair Hassan, Christopher C. Cline, Sara Parmigiani, Jade Truong, James W. Hartford, Nai-Feng Chen, Takako Fujioka, Scott Makeig, Alvaro Pascual-Leone, Corey J. Keller

**Author notes:** **Correspondence.** Jessica M. Ross, PhD, Stanford University, Department of Psychiatry and Behavioral Sciences, 401 Quarry Road, Stanford, CA 94305-5797.

## Abstract

Transcranial magnetic stimulation (TMS) applied to the motor cortex has revolutionized the study of motor physiology in humans. Despite this, TMS-evoked electrophysiological responses show significant variability, due in part to inconsistencies between TMS pulse timing and ongoing brain oscillations. Variable responses to TMS limit mechanistic insights and clinical efficacy, necessitating the development of methods to precisely coordinate the timing of TMS pulses to the phase of relevant oscillatory activity. We introduce Sensory Entrained TMS (seTMS), a novel approach that uses musical rhythms to synchronize brain oscillations and time TMS pulses to enhance cortical excitability. Focusing on the sensorimotor alpha rhythm, a neural oscillation associated with motor cortical inhibition, we examine whether rhythm-evoked sensorimotor alpha phase alignment affects primary motor cortical (M1) excitability in healthy young adults (*n*=33). We first confirmed using electroencephalography (EEG) that passive listening to musical rhythms desynchronizes inhibitory sensorimotor brain rhythms (*mu oscillations*) around 200 ms before auditory rhythmic events (27 participants). We then targeted this optimal time window by delivering single TMS pulses over M1 200 ms before rhythmic auditory events while recording motor-evoked potentials (MEPs; 19 participants), which resulted in significantly larger MEPs compared to standard single pulse TMS and an auditory control condition. Neither EEG measures during passive listening nor seTMS-induced MEP enhancement showed dependence on musical experience or training. These findings demonstrate that seTMS effectively enhances corticomotor excitability and establishes a practical, cost-effective method for optimizing non-invasive brain stimulation outcomes.

## 1. Introduction

Transcranial magnetic stimulation (TMS) is a widely used form of noninvasive brain stimulation with applications across basic and translational research and clinical medicine^1–3^. TMS is FDA-cleared for the treatment of depression, migraines, obsessive-compulsive disorder, smoking cessation, with more under investigation in Phase III clinical trials^4^. There is an accumulating literature on the effects of TMS on neurophysiology, cognition, behavior, and symptoms, but several systematic reviews and meta-analyses have revealed significant heterogeneity^5–11^ and low test-retest reliability^12–15^ in all domains of TMS research. In response to this challenge, efforts are being made to optimize TMS methods^7^.

One such approach to reducing variability of TMS effects is to employ brain state dependent neuromodulation. The targeted *brain states* in this context are times at which a brain network may be most sensitive to the effects of TMS^7,15–23^. Brain states can be quantified by analyzing endogenous brain oscillations as measured using electroencephalography (EEG). EEG studies demonstrate that the timing of TMS relative to these oscillations can significantly impact neural effects. Specifically, when TMS is applied to the primary motor cortex (M1) at specific phases of alpha frequency band activity, larger brain responses are evoked, as measured using motor evoked potentials (MEPs)^22–24^. Interacting with these phasic relationships, periods of *desynchronization* in endogenous sensorimotor mu oscillation (µ, activity recorded over somatomotor cortex with a fundamental in the alpha band) tend to coincide with longer timescale reductions in motor cortical excitability. Sensorimotor µ is associated with inhibitory control^25–27^, and its state of desynchronization correlates with cortical excitability. When µ is *desynchronized* cortical excitability is highest, and when µ is *synchronized* cortical excitability is lowest^28–30^. Together these findings suggest that applying TMS time-locked with periods of desynchronized µ (*i*.*e*., low phase alignment) may evoke larger brain responses^22,28^.

Leveraging this potential link between mu phase related cortical excitability and TMS related corticomotor excitability, studies now show that it is possible to enhance MEPs using µ in a way that is reliable^19^ and has been reproduced in multiple studies^19,21^. Moreover, repetitive TMS timed to these µ dynamics enhances changes in excitability (*i*.*e*., plasticity)^22^ and network changes across connected brain regions^19,21^. While these results are promising, EEG triggered TMS currently requires applying TMS pulses according to EEG recordings in real-time, making this technique difficult to implement in many research and clinical settings^18^. Even implementing EEG in clinic visits would require additional preparation time and resources including specialized staff. Further, the technique requires real-time signal processing with high temporal resolution, accurate EEG phase estimation algorithms, and closed-loop TMS-EEG systems. In a subset of individuals in whom a relevant and robust oscillatory signal cannot be measured, such EEG-triggered stimulation approaches can exhibit degradations in performance or fail entirely. Low cost and low resource alternative solutions are thus much needed to increase accessibility to phase-aligned TMS.

Outside of the TMS-EEG literature, there is an abundance of research showing that musical rhythms can reliably synchronize brain oscillations. Early work showed that music induces phase synchronization changes in beta and gamma bands in relation to musical beat times^31^. Since this work, beat-related phase alignments have been shown to be reproducible^32–34^, strongest for complex musical rhythms^33^, and present in multiple frequency bands including beta^31–37^, high beta/low gamma^38^, and alpha/µ^30^. This beat-related phase behavior is robust across stimuli and experimental designs^32,34,38^, modulates the connectivity between brain regions^32^, and reflects top-down aspects of perception^30,34,35,39–43^, and can be identified using *intertrial coherence* (*ITC*)^44^. Thus, *musical beats phase-align neural oscillations in multiple frequency bands*^30,35^ *and brain regions*^35^ *and this reflects dynamically shifting excitability brain states*^39–41^. These excitability dynamics around predictable musical beats should be relevant for corticomotor excitability when applying TMS to primary motor cortex. Stupacher *et al*. (2013)^45^ showed that music that induces more sensorimotor coupling can result in larger MEPs than music with less sensorimotor coupling, and that musical training can be relevant to this effect. This study provides a link between the literature on music-related sensorimotor dynamics and the TMS literature on corticospinal excitability, but the specific relationship between beat-related EEG dynamics and fluctuations in TMS excitability have yet to be investigated.

Here we introduce Sensory Entrained TMS (seTMS), which pairs auditory rhythms and TMS to align brain oscillations and enhance the effects of TMS. seTMS is a low cost and low resource alternative solution to EEG-triggered TMS that uses music to align the phase of relevant brain oscillations during TMS. Instead of timing TMS using real-time EEG recordings, rhythmic sensory events can be used to align the phase of cortical oscillations^46–52^ in preparation for TMS. By providing musical events around the TMS pulse, brain oscillations phase-shift to align with the musical beat events, and these shifts have a predictable timing relative to the musical events. Therefore, *one can predict the phase dynamics of excitability brain states using the musical event times alone without the need for EEG*. Synchronizing brain oscillations around the auditory beat enables the application of TMS pulses at the right time for maximal effect, when the phase of inhibitory oscillations are desynchronized, representing states of excitability. Using music to control phase alignment of brain waves during TMS has great potential to improve the neural effects of TMS in a low-cost, clinic-ready method.

In the current study we examine the effects of seTMS on corticomotor excitability (using the MEP). Specifically, we measured MEP sizes elicited after single pulses of seTMS compared to standard single pulse TMS to primary motor cortex. We hypothesized that seTMS, with TMS pulses timed with desynchronized inhibitory µ rhythms (high excitability state) driven by musical beats, would result in larger MEPs. Consistent with our hypothesis, we found that seTMS evoked larger MEPs compared with standard single pulse TMS. We also found larger MEPs when compared with an auditory control condition that used the same music but with alternate TMS timing. Years of musical experience or training did not significantly affect these results and thus this approach has the potential to substantially enhance TMS effects across all individuals. This work contributes to the growing understanding of interactions between brain oscillations and TMS and provides a low-cost and resource-efficient alternative for phase-aligned stimulation that may help address the heterogeneity of outcomes reported in TMS literature.

## 2. Methods

### 2.1. Participants and Study Design

This study was carried out in accordance with the Declaration of Helsinki. It was reviewed and approved by the Stanford University Institutional Review Board, performed in accordance with all relevant guidelines and regulations, and written informed consent was obtained from all participants. 37 healthy participants (22-65 years old [M=40.2, SD=14.6, 18F/18M/1O]) responded to an online recruitment ad and after an initial online screening and consent, 33 eligible participants (22-65 years old [M=39.8, SD=14.9, 17F/15M/1O]) were enrolled. Of the four who were not enrolled, two were excluded due to scheduling conflicts, one due to loss of interest, and one due to exclusion criteria. Of these, 20 enrolled for seTMS and 27 enrolled for EEG during listening to a rhythmic sound (with 14 participants enrolling for both seTMS and EEG during listening). In the end, *n*=27 participated in the EEG during listening. Of the 20 participants who enrolled for seTMS, one participant only participated in a subset of conditions, so the remaining *n*=19 participants were included in the MEP analyses. A total of *n*=13 participated in both EEG during listening and seTMS and were used in the analysis comparing EEG to MEP results. SeeTable 1 for *n*=33 demographics, and Supplementary Tables S1-3 for demographics of each study subgroup.

**Table 1.** Demographics. n=33

|  |  |
| --- | --- |
| Age, mean years (SD) | 39.8 (14.9) |
| Sex |  |
| Female, $n$ (%) | 17 (51.5) |
| Male, $n$ (%) | 15 (45.5) |
| Other or prefer not to state, $n$ (%) | 1 (3.0) |
| Handedness |  |
| Left hand dominant, $n$ (%) | 2 (6.1) |
| Right hand dominant, $n$ (%) | 31 (93.9) |
| Ambidextrous, $n$ (%) | 0 (0.0) |
| Education |  |
| GED or High School Diploma, $n$ (%) | 1 (3.0) |
| Some college, no degree, $n$ (%) | 2 (6.1) |
| Two year degree, $n$ (%) | 4 (12.1) |
| Four year degree, $n$ (%) | 16 (48.5) |
| Post graduate degree, $n$ (%) | 10 (30.3) |
| Employment |  |
| Part-time, $n$ (%) | 9 (27.3) |
| Full-time, $n$ (%) | 9 (27.3) |
| Unemployed, $n$ (%) | 9 (27.3) |
| Retired, $n$ (%) | 2 (6.1) |
| Part-time student, $n$ (%) | 1 (3.0) |
| Full-time student, $n$ (%) | 3 (9.1) |
| Race |  |
| White, $n$ (%) | 15 (45.5) |
| Black or African American, $n$ (%) | 4 (12.1) |
| American Indian or Alaska Native, $n$ (%) | 0 (0.0) |
| Asian, $n$ (%) | 11 (33.3) |
| Native Hawaiian or Other Pacific Islander, $n$ (%) | 1 (3.0) |
| Two or more races, $n$ (%) | 0 (0.0) |
| Some other race or prefer not to state, $n$ (%) | 2 (6.1) |

Inclusion criteria on the online screening form were (a) aged 18-65, (b) able to travel to study site, (c) fluent in English and (d) fully vaccinated against COVID-19. Exclusion criteria were (a) lifetime history of psychiatric or neurological disorder, (b) substance or alcohol abuse/dependence in the past month, (c) heart attack in the past 3 months, (d) pregnancy, (e) presence of any contraindications for TMS, such as history of epileptic seizures or certain metal implants^53^, or psychotropic medications that increase risk of seizures, and (f) Quick Inventory of Depressive Symptomatology (16-item, QIDS) self-report questionnaire score of 11 or higher indicating moderate depression^54,55^. All participants completed an MRI pre-examination screening form provided by the Richard M. Lucas Center for Imaging at Stanford University to ensure participant safety prior to entering the MRI scanner. Eligible participants were scheduled for two study visits: an anatomical MRI scan on the first visit and a TMS, EEG, or TMS with EEG session on the second visit.

### 2.2. Transcranial Magnetic Stimulation

#### TMS targeting and calibration

TMS was delivered using a MagVenture Cool-B65 A/P figure-of-eight coil from a MagPro X100 system (MagVenture, Denmark). TMS pulse triggering was automated to ensure correct timing in relation to the musical beats, using the MAGIC toolbox for MATLAB^56,57^. Neuronavigation (Localite TMS Navigator, Alpharetta, GA) using each participant’s MRI and a TMS-Cobot system (Axilum Robotics, France) were used to automatically maintain TMS coil placement relative to the subject’s head. MRI was performed on a GE DISCOVERY MR750 3T MR system (General Electric, Boston, Massachusetts) using a 32 channel head coil. T1 structural scans were acquired using a BRAVO pulse sequence (T1-weighted, sagittal slice thickness 1 mm, acquisition matrix 256 × 256, TR 8 ms, TE 3 ms, FA 15°).

### Resting motor threshold

To obtain resting motor threshold (RMT), single pulses of TMS were delivered to the hand region of the left primary motor cortex with the coil held tangentially to the scalp and at 45° from the midsagittal plane^58–60^. The optimal motor hotspot was defined as the coil position from which TMS produced the largest and most consistent MEP in a relaxed first dorsal interosseous (FDI) muscle^60^. RMT was determined to be the minimum intensity that elicited an MEP of at least 50 µV peak-to-peak amplitude in relaxed FDI in ≥ 5/10 stimulations^61,62^.

### Single pulse seTMS

Mu phase alignment dynamics occur around musical beat events and suggest that highest excitability (alpha desynchronization) may occur approximately 200 ms prior to the beat events^30,35^. To target this brain state with TMS, single pulses were applied at -200 ms in relation to the musical beat (Fig. 1). To assess whether seTMS increases excitability, we recorded MEPs in 20 participants that were evoked using standard single pulse TMS (hereafter referred to as standard TMS) and using single pulse seTMS (se-spTMS, hereafter referred to as seTMS), both applied for 100-150 trials at 120% of RMT. An additional auditory control condition was collected using the same auditory stimuli as used during seTMS but with TMS pulses applied at the same time as auditory beats (0 ms offset). Auditory stimuli were presented using earbuds at the maximum volume comfortable for each participant. These earbuds are also designed to be earplugs with a noise reduction rating (NRR) of 25 dB (Elgin USA Ruckus Earplug Earbuds, Arlington, Texas), intended to dampen the TMS “click” sound before reaching the ear canal. For additional dampening of the TMS “click” sound, we used over-the-ear noise-reducing foam-filled earmuffs (3M Ear Peltor Optime 105 behind-the-head earmuffs, NRR 29 dB, Maplewood, Minnesota). Our primary outcome measure was the MEP, averaged over the trials for each experimental condition. The order of TMS conditions was randomized across participants. We hypothesized that seTMS would evoke larger amplitude MEPs compared with standard TMS, even when using an auditory control.

**Fig. 1.**
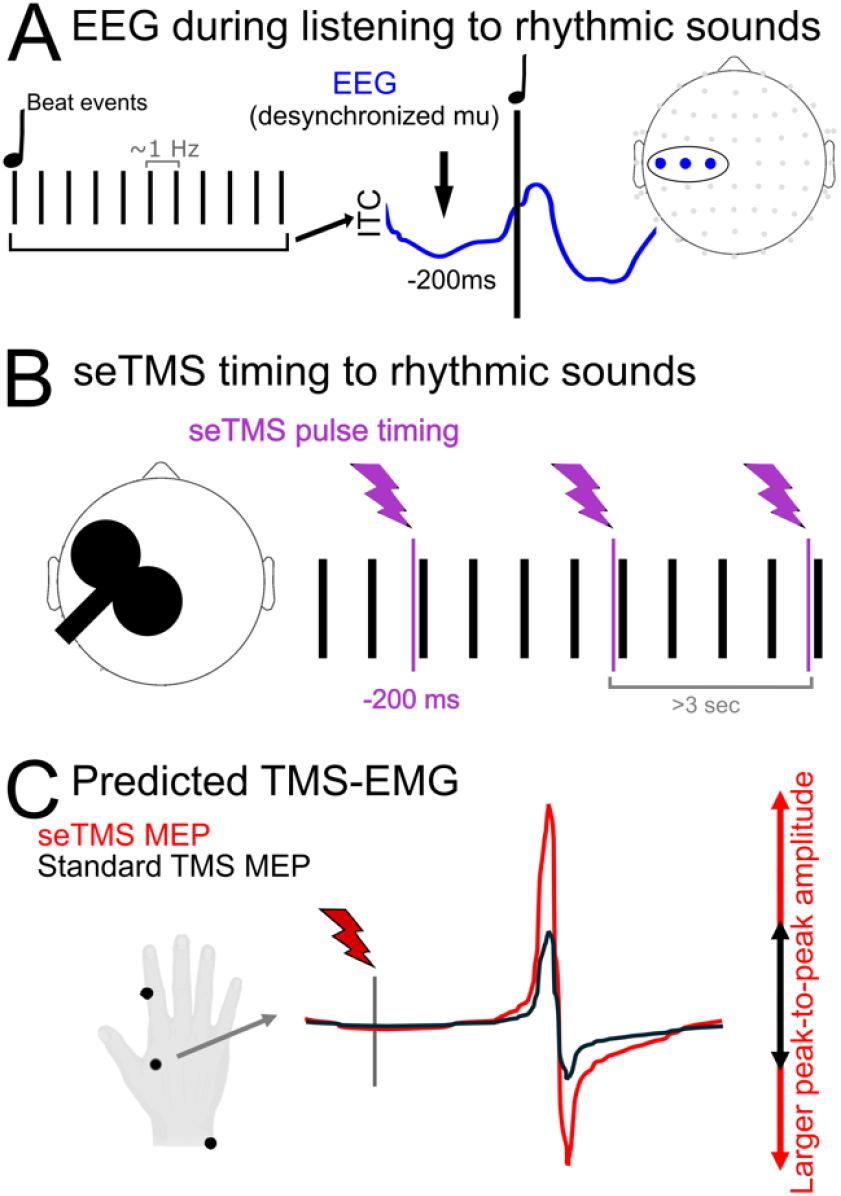
Study Design and seTMS Implementation. A) Desynchronization of mu occurs prior to beat events in musical rhythms and represents a high excitability state. Highest excitability states occur ∼200 ms prior to the musical beat events, regardless of musical tempo. B) TMS pulses were applied to the primary motor cortex using standard single pulse (standard spTMS) and single pulse seTMS (at 200 ms prior to musical beat events). C) Peak-to-peak amplitude of averaged motor-evoked potentials (MEPs) from EMG of the FDI muscle was used to assess excitability. Interstimulus interval lengths between TMS pulses were matched between standard TMS and seTMS conditions, and at least 3 seconds long. Musical sounds were played through earbud-earplugs and noise minimizing over-the-ear muffs were worn to reduce perception of TMS sounds.

### 2.3. Electromyography

Corticospinal excitability was measured using the peak-to-peak amplitude of motor evoked potentials (MEPs) recorded using electromyography (EMG) from the relaxed first dorsal interosseous (FDI) muscle of the right hand. One surface electrode was placed on the belly of the participants’ right FDI muscle. A reference electrode was placed on the lateral face of the proximal interphalangeal joint of the same finger as to not restrict movement. A ground electrode was placed on the styloid process of the wrist of the same hand. To obtain optimal EMG signal, the skin under the electrodes was abraded and cleaned and the electrodes were secured with medical tape. MEPs were elicited by applying single-pulse TMS to the region of the left motor cortex that induced MEPs in FDI. Participants were instructed to keep their head still and remain relaxed with their right hand on their lap for the duration of the experiment.

#### 2.3.1. Preprocessing of EMG

All collected EMG data were processed offline using customized automated scripts running in MATLAB. EMG data were baseline corrected by subtracting the mean value from 20 to 5 ms pre-TMS stimulation from the entire elicited signal. Next, trials with artifacts such as pre-activation or concurrent muscle activity were identified. To do this, the root mean square (RMS) of the EMG signal from -200 ms pre-TMS pulse to 13 ms post-TMS pulse, omitting -5 to +5 ms to avoid pulse artifact, was calculated. Trials with RMS values greater than 2.5 standard deviations (SD) from the average RMS of the entire block of trials were removed. Trials without a biphasic signal between 15 and 40 ms were excluded. Trials in which MEP amplitudes were larger than 5 standard deviations from the mean were excluded as outliers. The average number of MEP trials remaining after cleaning was 96.5 trials (SD = 21.8) for seTMS, 73.7 trials (SD = 18.7) for standard TMS, and 77.5 trials (SD = 16.1) for the auditory control condition.

### 2.4. Electroencephalography

In 27 participants, EEG was recorded during beat listening without TMS. This was for an individualized analysis of oscillatory phase-alignment within alpha and beta frequency bands. We expected that all participants would have music-induced excitability brain states. Further, we asked whether some aspects of musical experience would correlate with the strength of these excitability states. 64-channel EEG was obtained using a BrainVision actiCHamp Plus amplifier, with ActiCAP slim active electrodes in an extended 10–20 system montage (actiCHamp, Brain Products GmbH, Munich, Germany) with a 25 kHz sampling rate to reduce the spread of the pulse artifact^63^. EEG data were online referenced to Cz and recorded using BrainVision Recorder software v1.24.0001 (Brain Products GmbH, Germany). Impedances were monitored and percentage of channels with impedances <10 kΩ was 99.2 ±SD 2.4%. Electrode locations were digitized using Localite (Localite TMS Navigator, Alpharetta, GA).

#### 2.4.1. Preprocessing of EEG

EEG data were pre-processed offline using a custom designed Resting-state Semi-Automated Preprocessing pipeline (R-SAP, described below, available at https://github.com/jross4-stanford/R-SAP)^64^ and EEGLab v2021.1 in MATLAB R2021a (Mathworks, Natick, MA, USA). **R-SAP**. Data were epoched and downsampled to 1000 Hz. Low-pass (49 Hz) and high-pass (1 Hz) filters were applied using a zero-phase 4th order Butterworth filter. Conservative channel rejection and epoch rejection, and noise removal were applied using the *clean_rawdata* function (FlatlineCriterion = 5, ChannelCriterion = 0.8, BurstCriterion = 5, WindowCriterion = 0.5). Missing/removed channels were interpolated using spherical interpolation, and data were re-referenced to the average. The mean number of channels removed was 0.3 channels (SD = 0.7, range = 0-3). The mean number of epochs remaining was 96.6 epochs (SD = 8.8, range = 54-100). Because recordings were made with 64 channels, and the signals were unlikely to have that many independent sources, PCA was used to reduce dimensionality prior to ICA to 30 dimensions. This approach can improve decomposition^65,66^ and signal to noise ratio of large sources^67^. Fast independent component analysis (FastICA) was run^68^ and the Multiple Artifact Rejection Algorithm (MARA)^69,70^ was used to identify components with high likelihood of being non-brain artifacts (posterior_artifactprob > 0.30). These components were removed, and remaining components were reviewed using the open source TMS-EEG Signal Analyzer (TESA v1.1.0-beta) extension for EEGLAB^71,72^ (http://nigelrogasch.github.io/TESA/), allowing for additional components to be rejected by an expert reviewer if necessary. Mean number of components remaining after cleaning was 11.8 components (SD = 3.4, range = 6-18).

### 2.5. Auditory stimuli

Musical samples used for seTMS were duple or quadruple meter (even groupings of musical beats) and had a tempo of 98-120 beats per minute (BPM). Due to alternating strong and weak beat patterns, this tempo results in strong beats ∼once per second (1 Hz). We used three musical stimuli selected from the Groove Library (Table 2 for details) to ensure maximal predictive sensory and neural engagement with the musical beats^45,73–77^, each repeated five times. All auditory stimuli were 30 seconds in length with order randomized. For the EEG recording during listening, we used an 120 BPM auditory metronome with alternating strong and weak beat sounds (weak = 1/10 amplitude) that has been shown to induce the same excitability dynamics^35^. The auditory metronome consists of 262 Hz tones (middle C), with each tone lasting 60 ms and having a 10 ms duration rise and fall, generated using MATLAB. Like the music, the metronome has strong beats once per second.

**Table 2.** Musical Stimuli.

| Name | Artist | Groove Rating (0-127)* | Tempo (BPM) |
| --- | --- | --- | --- |
| Music | Leela James | 101.1 | 98 |
| Outa-Space | Billy Preston | 90.9 | 116 |
| Baby It's You | JoJo | 79.7 | 120 |

\*Note. Information taken from the Groove Library, compiled and rated by Janata *et al.* (2012)<sup>73</sup>.

### 2.6. Analyses

#### 2.6.1. Analysis of EEG

To observe oscillatory phase dynamics during beat listening, time-frequency analysis was completed for each participant at each channel. To focus on sensorimotor channels, the resulting time-frequency representations were then averaged across three channels from over the left motor cortex (C5, C3, C1). The time-frequency calculations were computed with the *newtimef* function in EEGLAB^78^ using linear spaced Morlet wavelets between 6 and 48 Hz with a fixed window size of 500 ms resulting in 3 cycles at the lowest frequency of 6 Hz. Log mean baseline power spectrum between 500 and 200 ms preceding beat times was removed^44,79,80^. The 500 ms window size was chosen to ensure that the time–frequency representation from each individual stimulus was not contaminated by either of the surrounding stimuli, which were 1000 ms apart. These computations were used to determine the event-related spectral perturbation (ERSP) in dB and phase coherence across trials (ITC)^78^. ITC is calculated by extracting the phase angle at each time–frequency point for each trial and comparing the phase angles across trials for coherence. This provides a coherence measure between 0 and 1, where 1 indicates complete coherence across trials for a given time–frequency point, and 0 indicates no coherence across trials.

Alpha activity was extracted from the ERSP values by averaging the power at each frequency bin between 8 and 14 Hz^26,27^. Alpha ITC was extracted using the same procedure except applied to ITC values instead of ERSP values. The same procedure was used to extract beta band ERSP and ITC between 20 and 26 Hz. Alpha ITC was used for the subsequent analyses on mu desynchronization dynamics. Troughs and peaks were calculated as the local minima and local maxima, between -222 and -99 ms and between 0 and 101 ms, respectively, for each individual participant. Oscillatory desynchronization followed by synchronization around an expected tone onset can be meaningfully represented by the slope, or the rise from ITC trough to ITC peak^35^. This measure is affected by both the amount and timing of ITC, and was calculated for all individual participants. Alpha ITC at trough versus at peak was compared using a paired sample *t*-test (*n*=27).

#### 2.6.2. Analysis of EMG

Peak-to-peak MEP amplitudes were calculated for the preprocessed EMG as the min-to-max voltage from 18 to 50 ms post-TMS. Percent change in MEP size between seTMS and standard TMS conditions was calculated using ((seTMS - standard TMS)/standard TMS))×100. MEP size was compared between conditions using a paired samples *t*-test (*n*=19). This percent change calculation and significance testing were then repeated to compare seTMS with the auditory control condition.

#### 2.6.3. Analysis of individual participant factors

We calculated the percentage of participants with larger MEPs in the seTMS condition, as well as the percent change in MEP size for these participants with an MEP gain. In order to explore whether having musical training or experience was associated with a participant’s exact alpha ITC trough time, we used an independent samples *t*-test to compare trough times across musicians and non-musicians in the 27 participants with EEG during music listening (*n*=14 musicians, *n*=13 non-musicians). Musicians were defined by having at least 1 year of musical training and/or experience (M = 7.93 years, SD = 4.93, range = 1 to 16). To explore whether years of musical experience or years *since* musical experience have a linear relationship with alpha ITC slope, we performed simple linear regression analyses. To explore whether being a musician resulted in a significant difference in percent change in MEP size, we performed an independent samples *t*-test using the 19 participants with MEP data (*n*=8 musicians, with M = 8.12 years of musical training and/or experience, SD = 6.47, range = 1 to 20). Lastly, to investigate whether there might be trends related to musicianship with regard to whether ITC at -200 ms or the time between ITC trough and -200 ms can predict MEP gain with seTMS, we used MEP data in all conditions and EEG during music listening from 13 participants (*n*=7 musicians, with M = 6.43 years of musical training and/or experience, SD = 4.68, range = 1 to 15) and plotted these variables against each other with a trend line. Although these groups are too small for a formal linear regression analysis, these exploratory investigations were intended to support future hypothesis generation about musician versus non-musician differences.

## 3. Results

### 3.1. Electroencephalography

To understand the effects of auditory beats on sensorimotor EEG, we first recorded EEG during beat listening without TMS and performed an individualized analysis of oscillatory phase-alignment within alpha and beta frequency bands. While participants listened to the auditory stimuli, EEG recorded over the motor cortex exhibited alpha frequency phase desynchronization (low coherence/ITC) and beta frequency phase synchronization (high coherence/ITC). This occurred in individual participants (Fig. 2A for a single participant and Fig. S1-2 for all individual participants) and in the group (Fig. 2B, *n*=27), reflecting a state of potentially increased motor excitability^25,26,30,35^. Music-induced phase dynamics showed an alpha ITC trough before each musical strong beat event (Fig. 2C, *n*=27, M = -156.48 ms, SD = 40.62) and an alpha ITC peak after the beat event (M = 46.41 ms, SD = 39.02), consistent with the literature^30,35,39^. These results are compatible with maximal motor excitability occurred ∼200 ms prior to musical beat events. ITC slope was positive in 26 out of 27 participants indicating that 96.30% of participants exhibited an alpha ITC desynchronization followed by a synchronization (Fig. 2C for all individual slopes). Alpha ITC was significantly smaller (Fig. 2D, *t*(26) = -8.34, *p* = 8.12×10^−9^) at the trough prior to the beat (M = 0.06, SD = 0.02) than at the peak after the beat (M = 0.11, SD = 0.03). For individual participant ITC and alpha ITC time series, see Supplementary Figs. S1-S2. Overall, these EEG findings during passive listening to musical rhythms confirm that we observed mu desynchronization around 200 ms before auditory rhythmic events.

**Fig. 2.**
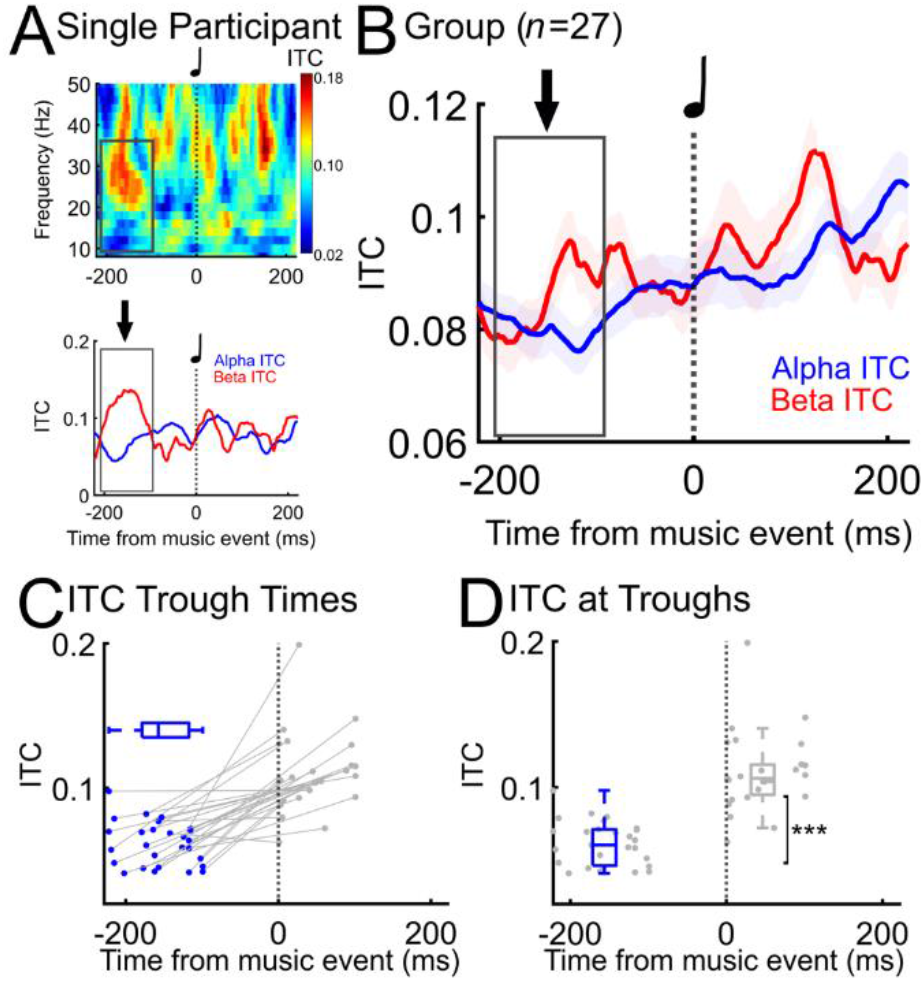
Auditory rhythms desynchronize mu. A) Individual participant music-induced motor cortex phase coherence in alpha (mu) and beta bands, with maximal excitability (low alpha/higher beta) occurring approximately 200 ms before beat events. Averaged across three channels from over the motor cortex (C5, C3, C1). B) Music-induced phase coherence in *n*=27 participants, with maximal excitability occurring approximately 200 ms before beat events. C) Individual participant (*n*=27) alpha ITC trough times (with box and whisker plot, alpha ITC peak times in gray, slopes from trough to peak in gray), and D) alpha ITC at trough vs. at peak (*** t(26) = -8.34, p = 8.12×10−9).

### 3.2. Electromyography

#### 3.2.1. Single pulse seTMS effects on the MEP

To target music-induced brain states with TMS, single pulses of TMS were applied to M1 at 200 ms prior to musical beat events (*i*.*e*., at the expected group ITC trough). One control condition was standard single pulse TMS without musical beats (referred to as *standard TMS*). Peak-to-peak MEP amplitudes were larger (*n*=19, Fig. 3 red vs. black, *t*(18) = 3.78, *p* = 0.0014) with seTMS ([M=3.08, SD=1.68, 95% CI=[2.27, 3.89]) compared with standard TMS ([M=2.44, SD=1.65, 95% CI=[1.64, 3.24]). The average percent increase in peak-to-peak amplitude from TMS to seTMS was 77.1% (median = 22.2%). An additional control condition used auditory beats with TMS pulses at 0 ms instead of at -200ms (referred to as *auditory control*). Peak-to-peak amplitudes were larger with seTMS (*n*=19, Fig. 3 red vs. gray, *t*(18) = 3.73, *p* = 0.0015) compared to the auditory control condition ([M=2.38, SD=1.56, 95% CI=[1.62, 3.12]). The average percent increase in peak-to-peak amplitude from the auditory matched condition to seTMS was 36.8% (median = 26.5). See Supplementary Figure S3 for all participants’ percent increase in MEP size, with group mean and median. These results suggest that seTMS enhanced corticomotor excitability over both standard TMS and an auditory control condition.

**Fig. 3.**
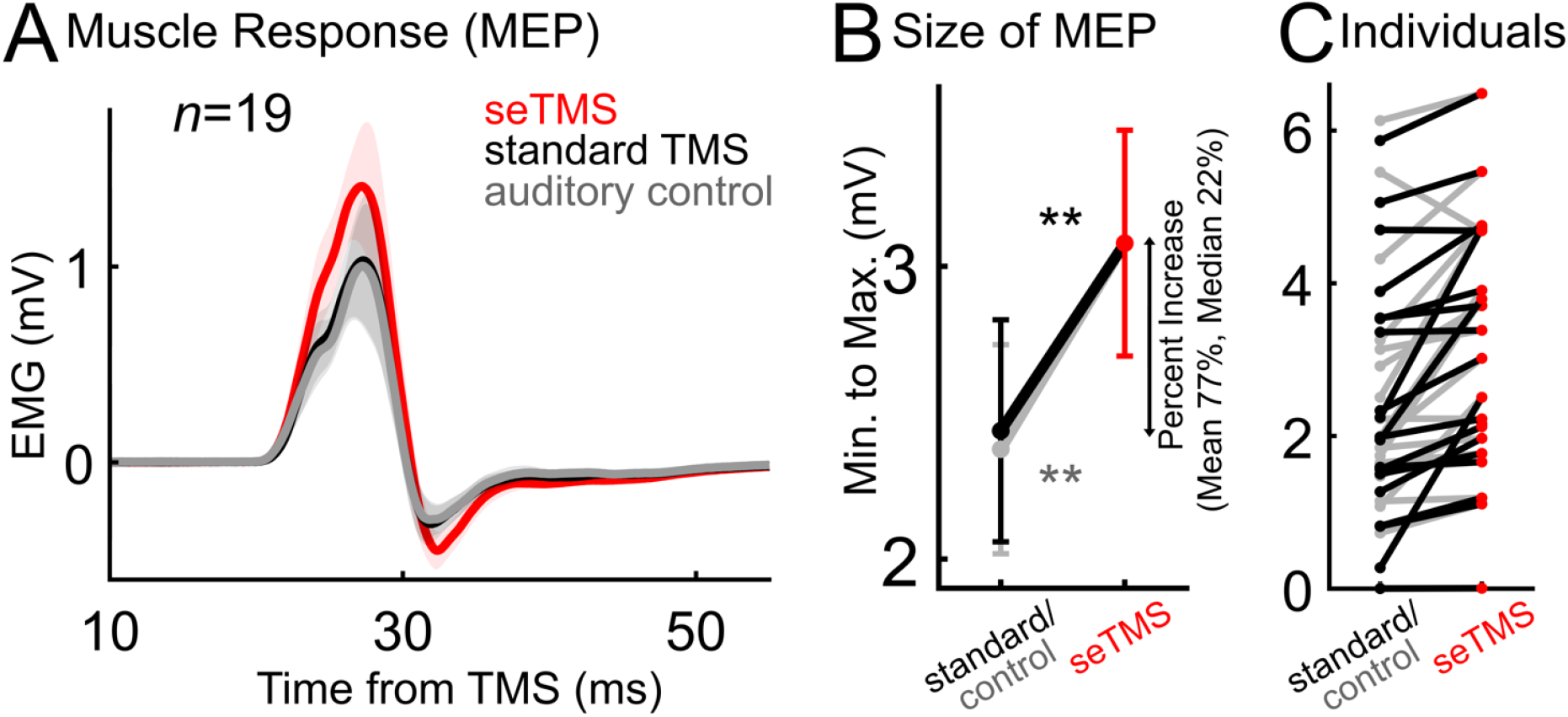
**seTMS increases the amplitude of motor-evoked potentials** compared with *standard TMS* and an *auditory control* condition. The auditory control condition used auditory matching to seTMS but with TMS pulses at 0 ms from the beat events. A) Motor-evoked potentials (MEPs) averaged over all participants (*n*=19). Shading represents standard error. B) Peak-to-peak amplitude mean (± standard error). Average percent increase from standard TMS = mean 77%, median 22%. (** black *t*(18) = 3.78, *p* = 0.0014; gray *t*(18) = 3.73, *p* = 0.0015). C) Individual participants.

### Individual Participant Factors

We next asked whether musical experience was relevant to individual participant seTMS effects on the MEP or to music-induced brain states. The MEP gain when using seTMS is present at the individual participant level in 18/19 of these participants (94.7%). Of the 18 participants with an MEP gain, the average percent increase was 81.4% but the percentage increase varied greatly across participants, ranging from <1% to >809%. We hypothesized that individual participant variability of the seTMS effect could be due to musical training or experience, which might affect how strong their phase dynamics are to the musical stimuli. In Table 3, experience and training is summarized for all participants. To explore whether having musical training or experience was associated with a participant’s exact alpha ITC trough time in the 27 participants with EEG during music listening, we compared trough times across musicians (at least 1 year of musical training and/or experience) and non-musicians with an independent samples *t*-test and found no difference between groups (*n*=27, Fig. 4A, *t*(25) = -0.39, *p* = 0.70). To explore whether years of musical experience or years since musical experience have a linear relationship with ITC slope, we performed simple linear regression analyses in the 27 participants with EEG during music listening and found the relationship to be non-significant (years *of* musical experience: *R*^2^ = 0.0030, *F*(1,26) = 0.076, *p* = 0.79; years *since* musical experience: *R*^2^=0.0071, *F*(1,26)=0.18, *p*=0.68). See also Fig. 4A for all individual participant and group average slopes and Fig. S3 for all individual ITC/slopes. Using the 19 participants with MEP data, we also found that having musical training or experience was not associated with a participant’s percent change in MEP size (*n*=19, Fig. 4B) when using seTMS compared with standard TMS (*t*(17) = 0.74, *p* = 0.47) or with the auditory control condition (*t*(17) = 0.88, *p* = 0.39). To explore whether years of musical experience or years since musical experience predicted percent change in MEP size, we performed simple linear regressions (*n*=19) and found that neither years of musical experience (compared with standard TMS: *R*^2^ = 0.01, *F*(1,11) = 0.15, *p* = 0.71; compared with auditory control: *R*^2^ = 0.02, *F*(1,11) = 0.23, *p* = 0.64) nor years since musical experience (compared with standard TMS: *R*^2^ = 0.08, *F*(1,11) = 0.95, *p* = 0.35; compared with auditory control: *R*^2^ = 0.003, *F*(1,11) = 0.03, *p* = 0.87) predicted percent change in MEP size. Using the 13 participants with both MEP data and EEG during music listening, we found similar relationships between ITC (at -200 ms, timing of ITC trough, time between ITC trough and -200ms) and percent change in MEP size when using seTMS compared with standard TMS or with the auditory control condition (*n*=13, Fig. S4). These overall null findings may suggest that seTMS is equally effective regardless of prior musical training or experience. Also see Supplementary Fig. S5-S8.

**Table 3.** Relevant training/experience. *n*=33 Non-musicians, *n* (%)

|  |  |
| --- | --- |
| Non-musicians, $n$ (%) | 18 (54.5) |
| Musicians, $n$ (%) | 15 (45.5) |
| Musical experience, mean years (SD, min-max) | 8.5 (5.8, 1-20) |
| Age experience began, mean years (SD, min-max) | 8.5 (3.2, 2.5-13) |
| Non-dancers, $n$ (%) | 25 (75.8) |
| Dancers, $n$ (%) | 8 (24.2) |
| Dance experience, mean years (SD, min-max) | 6.9 (8.5, 1-21) |
| Age experience began, mean years (SD, min-max) | 20.9 (18.1, 4-54) |
| No other physical hobbies, $n$ (%) | 17 (51.5) |
| Other physical hobbies, $n$ (%) | 16 (48.5) |
| Physical experience, mean years (SD, min-max) | 14.0 (10.5, 1-35) |
| Age experience began, mean years (SD, min-max) | 16.2 (10.6, 4-43) |

**Fig. 4.**
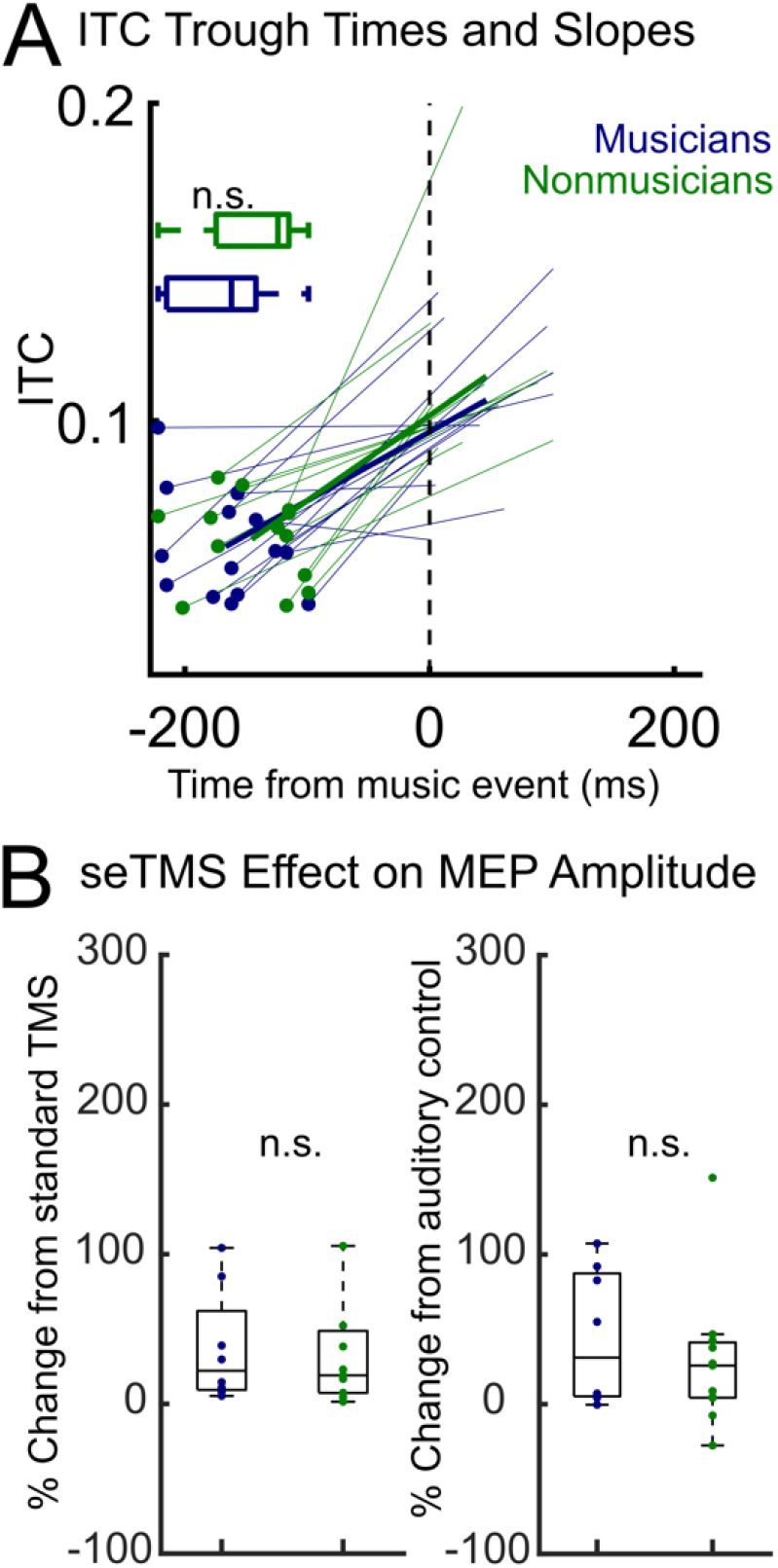
ITC dynamics and seTMS effects do not depend on musical experience. A) Individual participant ITC local minima (*t*(25) = -0.39, *p* = 0.70) and slopes (with group average slope shown using a thicker line) in musicians (*n*=14) and non-musicians (*n*=13). B) Individual participants’ increase in MEP size with seTMS in musicians (*n*=8) and nonmusicians (*n*=11), shown as percent change from standard TMS (*t*(17) = 0.74, *p* = 0.47) and from the auditory control condition (*t*(17) = 0.88, *p* = 0.39).

## 4. Discussion

In this study, we present a novel approach to TMS called Sensory Entrained TMS (seTMS) that uses music to synchronize excitability to prepare the brain for TMS (Figs. 1-2). We show that single pulses of seTMS to the primary motor cortex produce larger MEPs than conventional TMS (Fig. 3). To our knowledge, synchronizing excitability dynamics for TMS is a novel approach for maximizing stimulation effects. Because seTMS targets optimal *brain states* for TMS, it has the potential to enhance the effects of TMS in individuals, to contribute to efforts to reduce heterogeneity across the TMS literature, and to contribute to the growing understanding of interactions between brain oscillations and TMS. Unlike existing brain state methods that rely on EEG to estimate endogenous time windows during which the brain may be more sensitive to TMS, we use music to actively control the timing of optimal brain states for stimulation. This method is low resource and easy to implement in both research and clinical settings. We showed that single TMS pulses timed relative to musical beats evoke larger MEPs compared with an alternate timing and with standard TMS (Fig. 3). This study was designed using the literature on predictive sensorimotor dynamics during music listening but may have broad implications for noninvasive brain stimulation across basic and translational research and clinical medicine. However, more work is needed to fully understand music-induced excitability for use with TMS. Below we outline the relevant literature, limitations of the current work, and areas wherein future research is required.

### Neural mechanisms underlying seTMS

The excitability dynamics that occur around musical beats are thought to be related to timing prediction of sensory events^34,42,81^. Motor systems are known to be heavily involved while perceiving musical rhythms, as shown by imaging studies (^82^ for an analytic review). Moreover, EEG and MEG studies show coupling between sensory stimuli and neural oscillations that support body movement^30–35,83,84^. This phenomenon is often described as *covert action*^39,42,43,81,83,84^, occurring even in the absence of executed motor action^30–35^. Sound-synchronized movement must be planned for in advance, regardless of whether that movement is executed, and this motor planning appears to be the same for moving to or merely perceiving auditory rhythms^81,83,84^.

The reason for covert action is still being investigated, but theories that posit an essential role for accurate auditory perception^42,43,81,85^ are now supported by cases of impaired perception with disease-related^86–89^ or stimulation-induced^40,41,90,91^ brain lesions. Many theories exist to explain the relationship between sensory timing and covert action^81,85,92–96^, with an emerging understanding that this action-perception relationship is an actively predictive neural process^81,85,97,98^. Regardless of the reason for these excitability dynamics, their robust presence during passive music listening can be measured using MEG^31–34^ or EEG^30,35^ in numerous brain regions^30–35^. Using MEG, beat-related excitability dynamics have been reported in auditory and sensorimotor cortices and in the cerebellum, and the authors suggest that these recordings are the result of unexecuted auditory-motor coordination used for timing prediction^31–33^. Notably, these dynamics change to match when the beat times are predicted to occur, meaning that top-down influences on auditory perception drive the excitability dynamics^34^. Using EEG, beat-related excitability dynamics have been reported in premotor and motor networks^30^ as well as in the parietal, frontal, sensorimotor, and occipital cortices^35^.

These excitability dynamics around predictable musical beats should be relevant for corticomotor excitability when applying TMS to primary motor cortex^22,24,27,30,99^. Stupacher *et al*. (2013)^45^ demonstrated that this could be the case by measuring MEPs elicited with TMS time-locked with musical beats rated as high vs. low groove. Our data here show that TMS timed instead using mu phase-related excitability dynamics just prior to the beat increases the size of MEPs compared with on-beat and with standard TMS (Fig. 3). To understand interactions between groove and the seTMS effect, a comparison of high vs. low groove sounds using different mu phase relative timings for seTMS is needed.

### Selecting the most effective music for seTMS

There are several factors that can contribute to the degree of sensorimotor engagement and covert action with music; these include acoustic features^100^ such as RMS energy, RMS variability, pulse clarity “attack,” spectral flux, and low-frequency spectral flux^74^, as well as having the right amount of rhythmic syncopation^101^, complexity^77,101^, and beat salience^102–105^. However, these features can be selected for in aggregate by choosing music with a high groove rating. Groove is a well-studied psychological construct used to describe music and its relationship with sensorimotor entrainment^73,75,102,106,107^. High groove music spontaneously induces a sense of wanting to move^73,101^, increases spontaneous body movement^73,102^, increases coordinated and distributed muscle activity^77^, and improves sensorimotor synchronization to the beat^73^. Groove is consistently perceived and rated by musician and non-musician listeners, regardless of musical style^73,75,101,106,107^. Stupacher *et al*. (2013)^45^ showed that music that has a high groove ratings resulted in larger MEPs than music with low groove ratings. In the current study, we used high groove excerpts selected from the Groove Library to ensure maximal sensorimotor engagement^73^ (Table 2), but future work is needed to understand the relationship between this seTMS effect and differing levels of groove rating, specific acoustic features in music, and individual participant preferences or familiarity.

### The role of musicianship for enhanced neuromodulation with seTMS

Many studies show differences in the sensorimotor coupling and covert action depending on whether a person is a musician or a non-musician. These effects of musical training can be observed in spontaneous movement^102^ and muscle activity^77^ during high and low groove listening. Additionally, there may be a relationship between musical training and MEPs specifically^108–110^. Haueisen and Knösche (2001)^108^ found that pianists showed larger MEPs than nonpianists while listening to piano music. Rosenkranz *et al*. (2007)^109^ found that paired associative stimulation combined with TMS had a larger effect on MEP size in musicians as compared to non-musicians. Stupacher *et al*. (2013) also showed that having musical training can be relevant to an MEP effect^45^. In a study looking specifically at plasticity induction, Kweon *et al*. (2023) found that 10 Hz rTMS paired with an NMDA receptor partial agonist increased MEP size in musicians and athletes more so than in non-musicians and non-athletes^111^. These results may be indicative of a direct relationship between musical or general motor skill training and increased synaptic connectivity and plasticity, a higher gain in cortical output, and/or more automated motor programming processes. However, some reports suggest no differences between MEPs in musicians and non-musicians^110,111^. Further, there appears to be individual variability in sensorimotor synchronization that is unrelated to musical training or experience, and has been suggested to be better explained by differences in beat extraction^112^. This may include varying functionality in brain structures involved in time perception and action integration or differences in strategy unrelated to training. Our results did not reveal any significant differences between MEPs or ITC factors in these two groups (Figs. 4, S3-8), necessitating more research to untangle individual variability and which training factors may be relevant. While null results indicate the potential for seTMS to be more widely effective, we suggest that the effects of musical training on both MEPs and on synchronized excitability with music should still be explored further to determine any potential relevance to seTMS personalization.

### Brain networks for enhanced neuromodulation with seTMS

The networks of the brain where we see covert action during music listening vary. Brain imaging during rhythm perception experiments consistently show activation in areas of the brain that are known to be involved in movement of the body, including primary motor cortex, premotor cortices, the basal ganglia, posterior parietal cortex, supplementary motor area, and cerebellum. A recent ALE (Activation Likelihood Estimation^113^) meta-analysis across 42 PET and fMRI studies of passive music listening investigated which activations were common across studies^82^. This analysis revealed that the premotor cortex, primary motor cortex, and a region of left cerebellum were most reliably and consistently implicated across studies. Interestingly, the authors also showed that stimulus variability across studies (such as acoustic features, instructions on how to attend to the music, emotional states, arousal, familiarity, attention and memory) did not have clear impacts on whether covert action was reported but only on which motor networks were covertly activated. Using MEG and EEG, beat-related excitability dynamics have been reported in sensory^31–35^, premotor^30,35^, motor^30–35^, frontal and parietal networks^30,35^. The integration of intracranial EEG (iEEG) and single-cell recordings could significantly enhance the localization of ITC effects, thereby maximizing the efficacy of seTMS. These techniques offer more localized and high spatiotemporal resolution compared with conventional EEG alone. Further, combining seTMS with iEEG to measure intracranial TMS evoked potentials (iTEPs) could provide deeper insights into neural mechanisms at the level of local circuit dynamics and trans-synaptic plasticity^114^. This approach may yield valuable knowledge about the causal relationships between sensory entrainment, connectivity patterns, and cognitive processes. Here we targeted the primary motor cortex because of the clear link with covert action and mu dynamics and because TMS to M1 provides a robust read-out in the MEP. However, future work should explore whether stimulation effects can be improved with music when applied to other brain targets, including nodes of implicated motor networks in covert action during music listening^82^, dorsal auditory stream^40,85^, and fronto-striatal pathways^115,116^.

### Translation to clinical practice

seTMS has the potential to substantially enhance the effects of TMS. Since seTMS does not require EEG, it is affordable and accessible, and could be quickly and easily adopted for clinical use. However, for seTMS to be relevant for psychiatric applications of TMS, it will be necessary to determine whether seTMS enhances the TMS-evoked EEG responses when applied to the dorsolateral prefrontal cortex (dlPFC), the treatment target for most psychiatric conditions treated with TMS. Due to beat-related excitability dynamics outside of motor cortex, including in fronto-striatal pathways sensitive to TMS^115,116^, seTMS may be relevant for dlPFC brain networks. Clinical TMS with concurrent music listening has been shown to be feasible and also effective for treating depression^117^, but using music to create excitability states for optimized treatment protocols has not previously been done.

## 5. Limitations and future directions

While our study demonstrates the potential of seTMS to enhance motor cortex excitability, future work should evaluate whether this approach can be used to induce plasticity. Several limitations should be addressed in future research. First, we focused solely on the primary motor cortex; future studies should explore the effects of seTMS on other brain regions, particularly the dorsolateral prefrontal cortex, given its relevance in treating psychiatric conditions. Second, our study did not include a clinical population, limiting our ability to draw conclusions about therapeutic potential. Third, we used a standardized set of musical stimuli; future work should investigate personalized music selection to optimize individual responses. Moving forward, key directions for research include: 1) developing repetitive seTMS protocols to induce lasting plasticity, 2) investigating seTMS effects in other brain regions, particularly those relevant to mood and emotion regulation, 3) exploring the potential for personalization of seTMS parameters, including music selection and timing, and 4) examining seTMS effects on cognitive tasks and in clinical populations.

## 6. Conclusions

In this study, we introduced Sensory Entrained Transcranial Magnetic Stimulation (seTMS), a novel approach that leverages music-induced changes in neural oscillations to enhance the effects of TMS. We demonstrated that seTMS significantly increased the size of motor-evoked potentials compared to standard TMS and an auditory control condition, with an average MEP increase of 77%. These effects were observed across participants, regardless of musical experience. By synchronizing TMS pulses with music-induced high-excitability brain states, seTMS offers a low-cost, accessible method to potentially reduce intra- and inter-individual variability in TMS responses. This approach opens new avenues for optimizing non-invasive brain stimulation techniques and may have significant implications for both research and clinical applications of TMS.

## Acknowledgments

We would like to acknowledge the contributions of all of our research participants. We extend gratitude to the members of the Precision Neurotherapeutics Laboratory for helpful feedback on the article and throughout the course of the study.

## Author contributions

JMR, JG, UH, CCC, SP, TF, SM, APL, and CJK conceptualized and designed the study. JMR and CJK acquired funding. JMR and JT programmed the experiment. JMR, JT, LF, and JWH collected the data. JMR and LF conducted the analyses. All authors interpreted the results. All authors contributed to the writing of the manuscript. All authors provided intellectual contributions to and approval of the final manuscript.

## Availability of Data and Materials

The datasets generated and/or analyzed during the current study are available upon request.

## Supplementary Information

The online version contains supplementary material.

