## Supplementary for "Sensory Entrained TMS (seTMS) enhances motor cortex excitability"

***Table S1. Demographics.*** *n*=27

| | |
|---|---|
| Age, mean years (SD) | 39.6 (14.2) |
| Sex | |
| Female, *n* (%) | 14 (51.9) |
| Male, *n* (%) | 12 (44.4) |
| Other or prefer not to state, *n* (%) | 1 (3.7) |
| Handedness | |
| Left hand dominant, *n* (%) | 2 (7.4) |
| Right hand dominant, *n* (%) | 25 (92.6) |
| Ambidextrous, *n* (%) | 0 (0.0) |
| Education | |
| GED or High School Diploma, *n* (%) | 1 (3.7) |
| Some college, no degree, *n* (%) | 2 (7.4) |
| Two year degree, *n* (%) | 4 (14.8) |
| Four year degree, *n* (%) | 13 (48.2) |
| Post graduate degree, *n* (%) | 7 (25.9) |
| Employment | |
| Part-time, *n* (%) | 9 (33.3) |
| Full-time, *n* (%) | 7 (25.9) |
| Unemployed, *n* (%) | 8 (29.6) |
| Retired, *n* (%) | 1 (3.7) |
| Part-time student, *n* (%) | 0 (0.0) |
| Full-time student, *n* (%) | 2 (7.4) |
| Race | |
| White, *n* (%) | 12 (44.4) |
| Black or African American, *n* (%) | 4 (14.8) |
| American Indian or Alaska Native, *n* (%) | 0 (0.0) |
| Asian, *n* (%) | 9 (33.3) |
| Native Hawaiian or Other Pacific Islander, *n* (%) | 1 (3.7) |
| Two or more races, *n* (%) | 0 (0.0) |
| Some other race or prefer not to state, *n* (%) | 1 (3.7) |

***Table S2. Demographics.*** *n*=19

| | |
|---|---|
| Age, mean years (SD) | 37.7 (14.3) |
| Sex | |
| Female, *n* (%) | 8 (42.1) |
| Male, *n* (%) | 10 (52.6) |
| Other or prefer not to state, *n* (%) | 1 (5.3) |
| Handedness | |
| Left hand dominant, *n* (%) | 1 (5.3) |
| Right hand dominant, *n* (%) | 18 (94.7) |
| Ambidextrous, *n* (%) | 0 (0.0) |
| Education | |
| GED or High School Diploma, *n* (%) | 0 (0.0) |
| Some college, no degree, *n* (%) | 2 (10.5) |
| Two year degree, *n* (%) | 2 (10.5) |
| Four year degree, *n* (%) | 10 (42.1) |
| Post graduate degree, *n* (%) | 5 (10.5) |
| Employment | |
| Part-time, *n* (%) | 4 (21.0) |
| Full-time, *n* (%) | 8 (42.1) |
| Unemployed, *n* (%) | 4 (21.0) |
| Retired, *n* (%) | 1 (5.3) |
| Part-time student, *n* (%) | 1 (5.3) |
| Full-time student, *n* (%) | 1 (5.3) |
| Race | |
| White, *n* (%) | 8 (42.1) |
| Black or African American, *n* (%) | 2 (10.5) |
| American Indian or Alaska Native, *n* (%) | 0 (0.0) |
| Asian, *n* (%) | 7 (36.8) |
| Native Hawaiian or Other Pacific Islander, *n* (%) | 1 (5.3) |
| Two or more races, *n* (%) | 0 (0.0) |
| Some other race or prefer not to state, *n* (%) | 1 (5.3) |

***Table S3. Demographics.*** *n*=13

| | |
|---|---|
| Age, mean years (SD) | 36.1 (12.0) |
| Sex | |
| Female, *n* (%) | 5 (38.5) |
| Male, *n* (%) | 7 (53.8) |
| Other or prefer not to state, *n* (%) | 1 (7.7) |
| Handedness | |
| Left hand dominant, *n* (%) | 1 (7.7) |
| Right hand dominant, *n* (%) | 12 (92.3) |
| Ambidextrous, *n* (%) | 0 (0.0) |
| Education | |
| GED or High School Diploma, *n* (%) | 0 (0.0) |
| Some college, no degree, *n* (%) | 2 (15.4) |
| Two year degree, *n* (%) | 2 (15.4) |
| Four year degree, *n* (%) | 7 (53.8) |
| Post graduate degree, *n* (%) | 2 (15.4) |
| Employment | |
| Part-time, *n* (%) | 4 (30.8) |
| Full-time, *n* (%) | 6 (46.2) |
| Unemployed, *n* (%) | 3 (23.1) |
| Retired, *n* (%) | 0 (0.0) |
| Part-time student, *n* (%) | 0 (0.0) |
| Full-time student, *n* (%) | 0 (0.0) |
| Race | |
| White, *n* (%) | 5 (38.5) |
| Black or African American, *n* (%) | 2 (15.4) |
| American Indian or Alaska Native, *n* (%) | 0 (0.0) |
| Asian, *n* (%) | 5 (38.5) |
| Native Hawaiian or Other Pacific Islander, *n* (%) | 1 (7.7) |
| Two or more races, *n* (%) | 0 (0.0) |
| Some other race or prefer not to state, *n* (%) | 0 (0.0) |

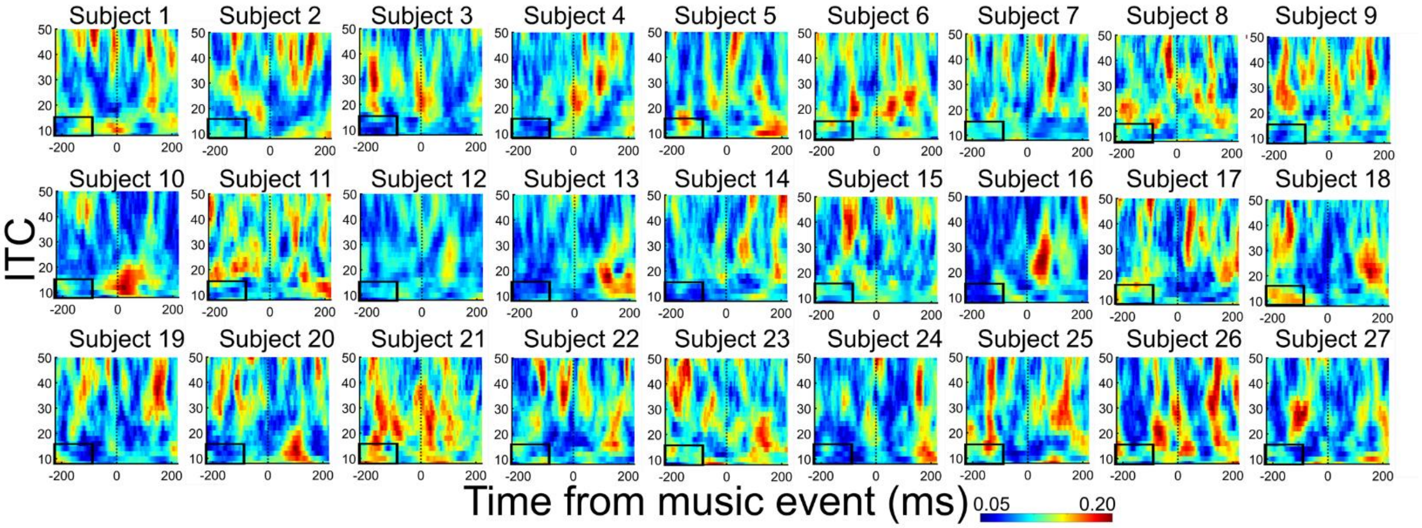


**Fig. S1. Individual participant ITC**, averaged across three channels from over the motor cortex (C5, C3, C1) and with frequency/time window ROI indicated with a black box (*n*=27)**.**

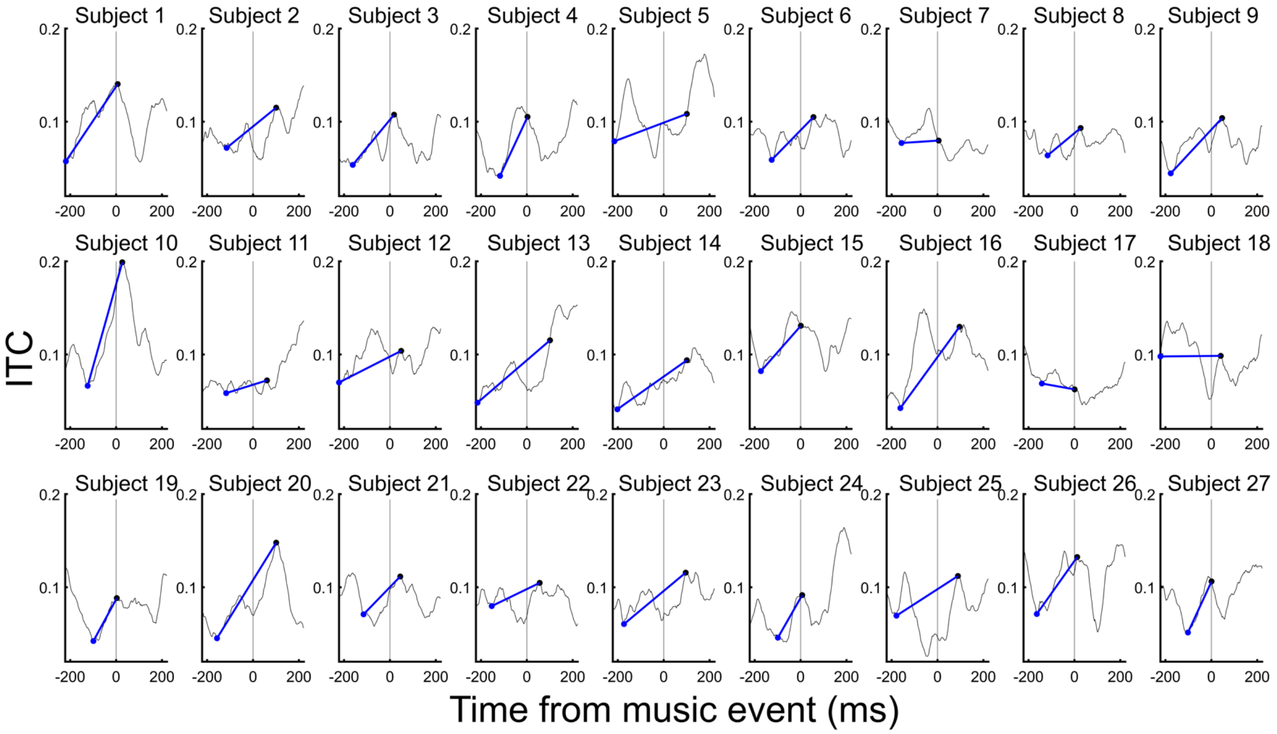


**Fig. S2. Individual participant ITC dynamics.** Shown are alpha ITC with local minima and maxima and slope (*n*=27).

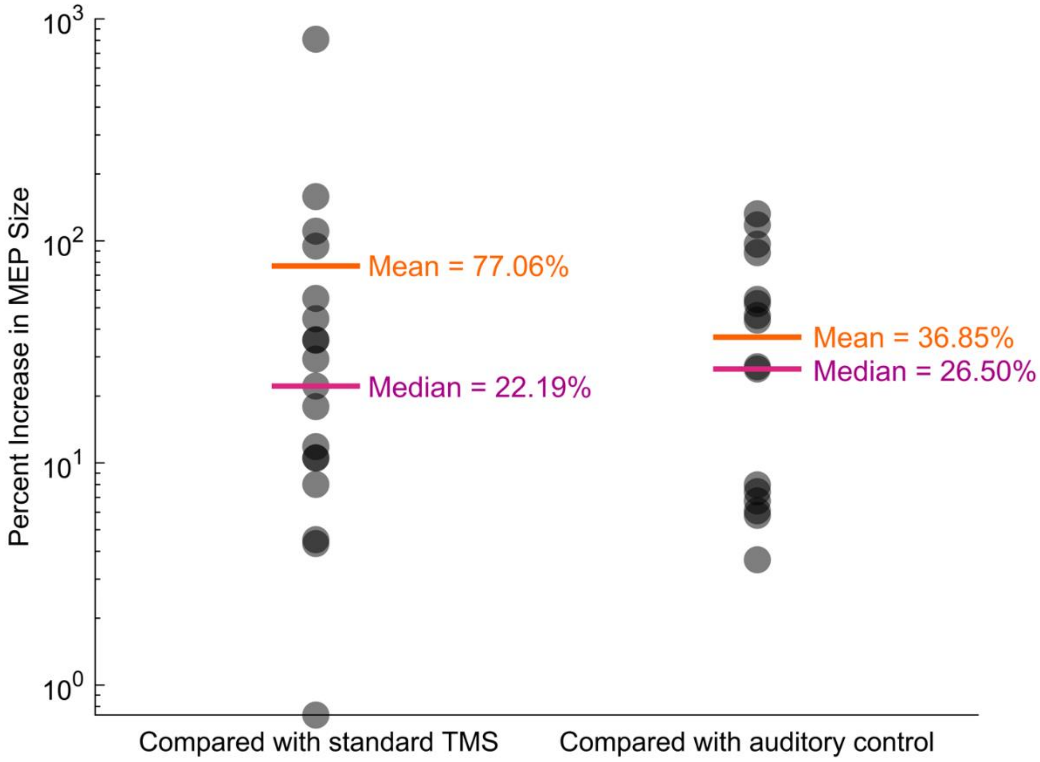


**Fig. S3. Percent increase in MEP size with seTMS**, compared to standard TMS (left) and to the auditory control (right). Mean and median percent increase is shown in orange and magenta, respectively (*n*=19).

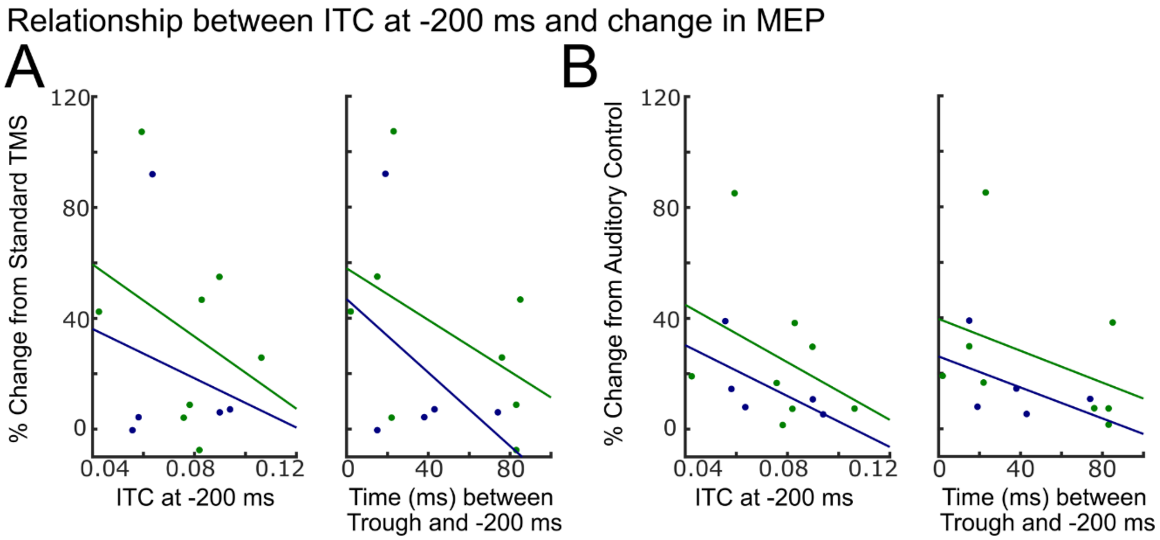


**Fig. S4. Relationship between ITC and increase in MEP size, in musicians and nonmusicians** (*n*=13). Results shown are using ITC at -200 ms and using individual participant ITC trough time, and increase in MEP size was calculated using the A) standard TMS and B) auditory control conditions.

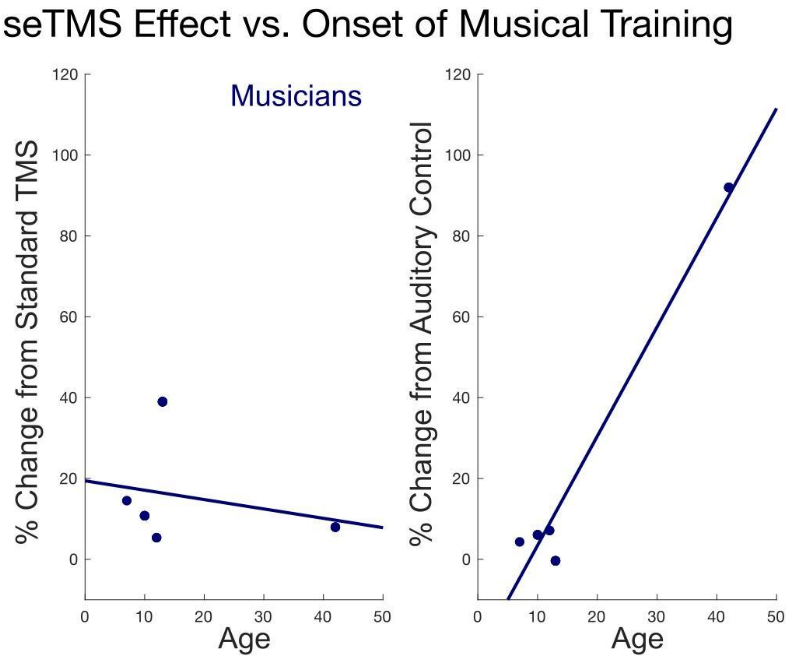


**Fig. S5. seTMS effect by age at which musical training began amongst musicians (*n*=5).**

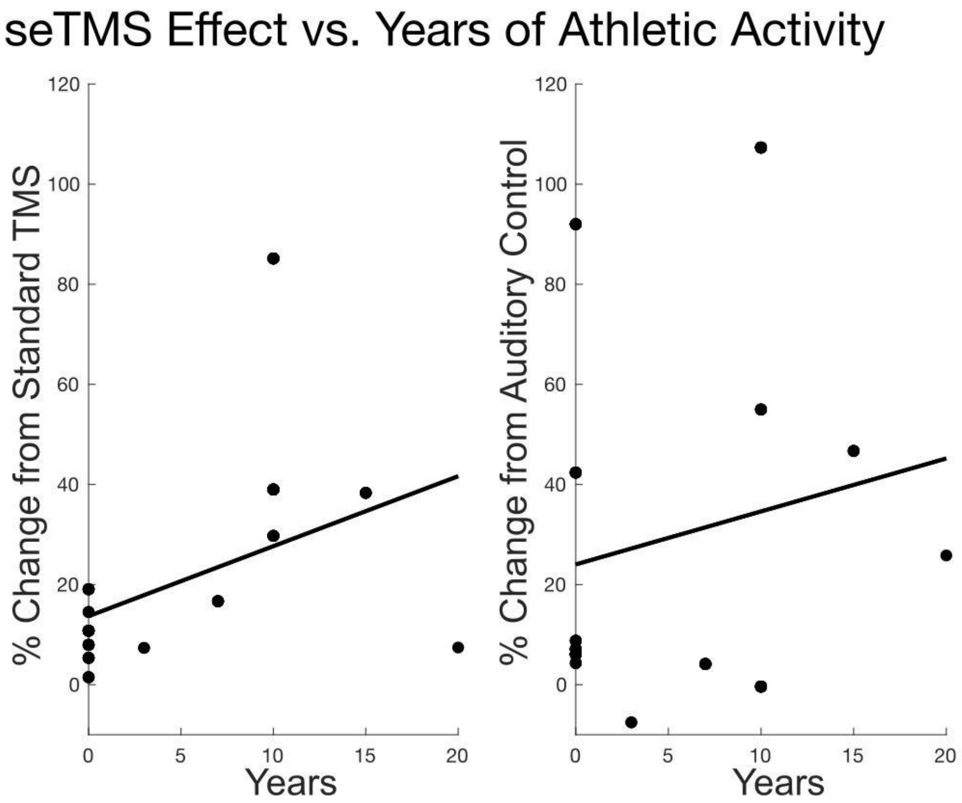

**Fig. S6. seTMS effect for all subjects by years of athletic activity (*n*=13).**

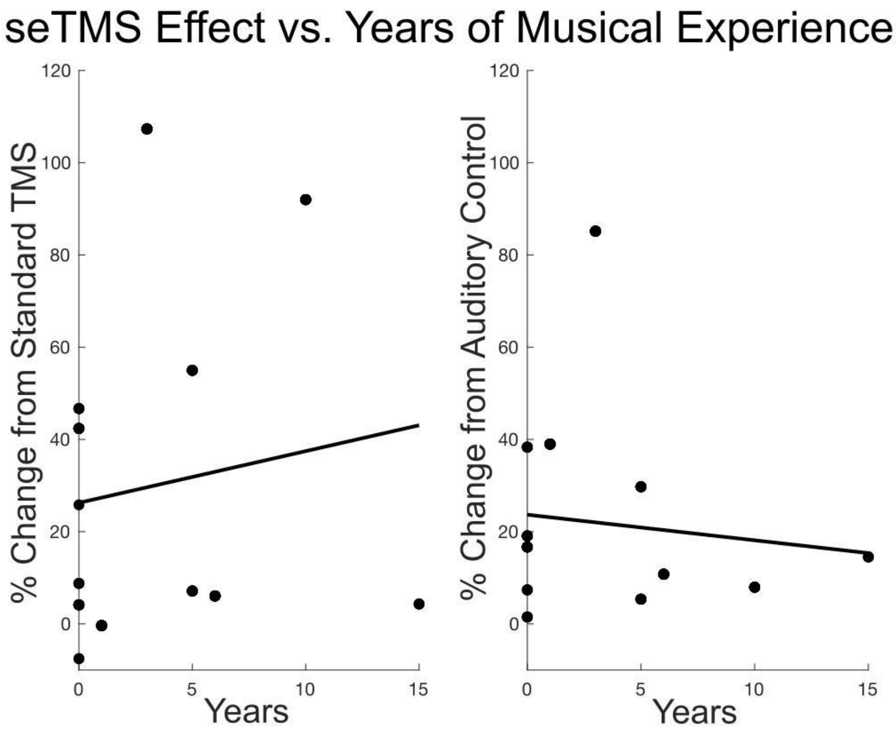


**Fig. S7. seTMS effect for all subjects by years of musical training or experience (*n*=13).**

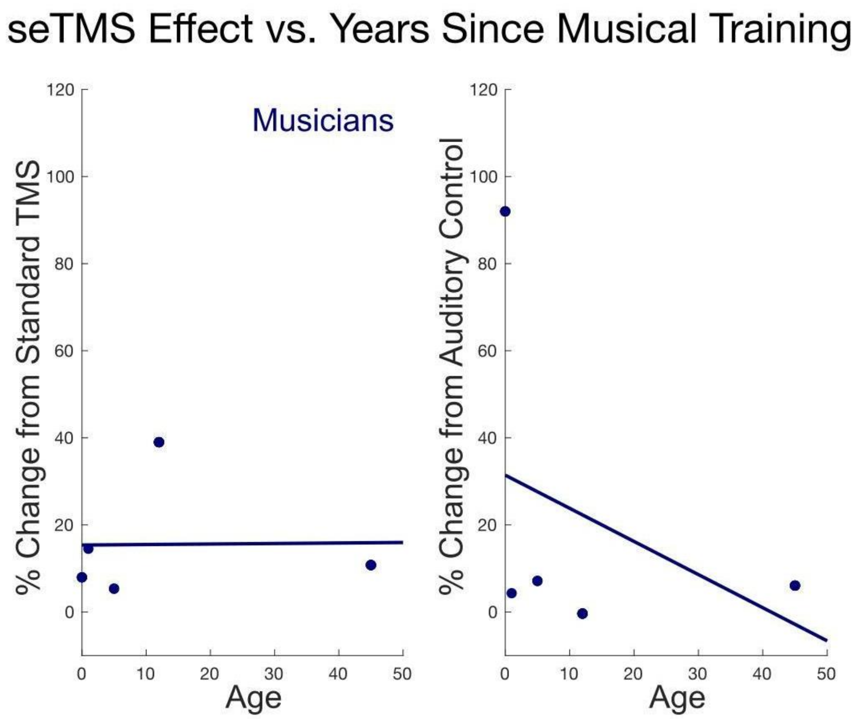


55 **Fig. S8. seTMS effect by years since most recent musical training or experience amongst**
56 **musicians (*n*=5).**
